# A mechanism for how Cdr1/Nim1 kinase promotes mitotic entry by inhibiting Wee1

**DOI:** 10.1101/730242

**Authors:** Hannah E. Opalko, Isha Nasa, Arminja N. Kettenbach, James B. Moseley

## Abstract

To enter into mitosis, cells must shut off the cell cycle inhibitor Wee1. SAD family protein kinases regulate Wee1 signaling in yeast and humans. In *S. pombe*, two SAD kinases (Cdr1/Nim1 and Cdr2) act as upstream inhibitors of Wee1. Previous studies found that *S. pombe* Cdr1/Nim1 directly phosphorylates and inhibits Wee1 *in vitro*, but different results were obtained for budding yeast and human SAD kinases. Without a full understanding of Cdr1 action on Wee1, it has been difficult to assess the *in vivo* relevance and conservation of this mechanism. Here, we show that both Cdr1 and Cdr2 promote Wee1 phosphorylation in cells, but only Cdr1 inhibits Wee1 kinase activity. Inhibition occurs when Cdr1 phosphorylates a cluster of serine residues linking α-helices G and H of the Wee1 kinase domain. This region is highly divergent among different Wee1 proteins, consistent with distinct regulatory mechanisms. A *wee(4A)* mutant that impairs phosphorylation by Cdr1 delays mitotic entry and causes elongated cells. By disrupting and re-targeting Cdr1 localization, we show that Cdr1 inhibition of Wee1 occurs in cells at cortical nodes formed by Cdr2. Based on our results, we propose a two-step model for inhibition of Wee1 by Cdr1 and Cdr2 at nodes.

## Introduction

Eukaryotic cells enter into mitosis due to regulated activation of Cdk1. During interphase, Cdk1 is kept inactive by the protein kinase Wee1, which phosphorylates Cdk1-Y15 to inhibit Cdk1 activity (Nurse, 1975; Gould and Nurse, 1989; Featherstone and Russell, 1991; Lundgren *et al*., 1991; Parker *et al*., 1992; Coleman *et al*., 1993). As cells enter mitosis, this inhibitory phosphorylation is removed by the phosphatase Cdc25 to activate Cdk1 (Russell and Nurse, 1986; Dunphy and Kumagai, 1991; Gautier *et al*., 1991; Kumagai and Dunphy, 1991; Strausfeld *et al*., 1991). To enter mitosis, cells must inhibit Wee1, thereby relieving the “brake” on Cdk1. This mechanism acts as a bistable switch due to feedback in which Cdk1 inhibits Wee1 and activates Cdc25 (Mueller *et al*., 1995; Pomerening *et al*., 2003; Sha *et al*., 2003; Harvey *et al*., 2005; Kim and Ferrell, 2007). Many upstream mechanisms that regulate Wee1 are not well defined. The fission yeast *S. pombe* has served as a long-standing model system for this conserved regulatory module. These rod-shaped cells enter into mitosis and divide at a reproducible size due to the activities of Wee1, Cdc25, and other Cdk1 regulators. Decades of work identified key factors upstream of Cdk1, but it has remained a challenge to place these factors into defined pathways and to understand their biochemical mechanisms.

Genetic screens in fission yeast defined two SAD-family (<u>S</u>ynapses of the <u>A</u>mphid <u>D</u>efective) protein kinases, Cdr1/Nim1 and Cdr2, as upstream inhibitors of Wee1. Both *cdr1* and *cdr2* mutants divide at a larger size than wild type cells due to uninhibited Wee1 (Russell and Nurse, 1987; Young and Fantes, 1987; Breeding *et al*., 1998; Kanoh and Russell, 1998). The cell size defects of *cdr1* and *cdr2* mutants are non-additive (Feilotter *et al*., 1991; Martin and Berthelot-Grosjean, 2009; Moseley *et al*., 2009), suggesting redundant or related inhibitory mechanisms. Wee1 becomes increasingly phosphorylated as cells grow during G2 (Lucena *et al*., 2017). This phosphorylation is reduced in *cdr1*Δ and *cdr2*Δ mutants (Allard *et al*., 2018), consistent with increasing inhibition by Cdr1-Cdr2. Cdr2 appears to act primarily through localization of Wee1. Cdr2 forms cortical node structures in the cell middle, and recruits both Cdr1 and Wee1 to these sites (Morrell *et al*., 2004; Martin and Berthelot-Grosjean, 2009; Moseley *et al*., 2009; Allard *et al*., 2018). Cdr1/Nim1 appears to be the key protein that directly regulates Wee1 activity. *In vitro*, Cdr1 phosphorylates and inhibits Wee1 kinase activity (Coleman *et al*., 1993; Parker *et al*., 1993; Wu and Russell, 1993). However, key questions have remained open regarding this mechanism. The *in vivo* relevance of Cdr1 phosphorylating Wee1 has not been tested because direct phosphorylation sites have been unknown. In addition, different results have been obtained for regulation of Wee1 by SAD family kinases in budding yeast and humans (Kellogg, 2003; Lu *et al*., 2004; Sakchaisri *et al*., 2004; Keaton and Lew, 2006). In this study, we investigated how Cdr1 inhibits Wee1. Our combined results lead to a mechanistic model for Wee1 regulation by SAD kinases in fission yeast.

## Results and Discussion

### Cdr1 phosphorylates and inhibits Wee1

Cdr1 and Cdr2 act as upstream inhibitors of Wee1 (Fig 1A). To test their mechanisms, we overexpressed Cdr1, Cdr2, or the empty vector in *cdr1*Δ*cdr2*Δ cells. We monitored Wee1 phosphorylation by SDS-PAGE band shift (Lucena *et al*., 2017; Allard *et al*., 2018), and Wee1 activity by Cdk1-pY15 levels (Fig 1B). Cdr1 overexpression induced hyperphosphorylation of Wee1 and loss of Cdk1-pY15, indicating inhibition of Wee1 kinase activity, consistent with previous work (Coleman *et al*., 1993; Parker *et al*., 1993; Wu and Russell, 1993). In contrast, Cdr2 overexpression induced hyperphosphorylation of Wee1 but no change in Cdk1-pY15. Thus, both Cdr1 and Cdr2 induce phosphorylation of Wee1, but the regulatory output of these two kinases is distinct. Consistent with these biochemical results, only over-expression of Cdr1 but not of Cdr2 resulted in reduced cell size in *cdr1*Δ*cdr2*Δ cells (Fig 1C), consistent with previous results in wild type cells (Russell and Nurse, 1987; Breeding *et al*., 1998). Phosphorylation of Wee1 in fission yeast cells was reduced in the catalytically inactive mutant *cdr1(K41A)* (Fig 1D), similar to *cdr1*Δ cells (Allard *et al*., 2018). Given the role of Cdr1 in regulating Wee1 kinase activity, along with open questions regarding its underlying mechanism, we investigated this pathway further.

**Figure 1:**
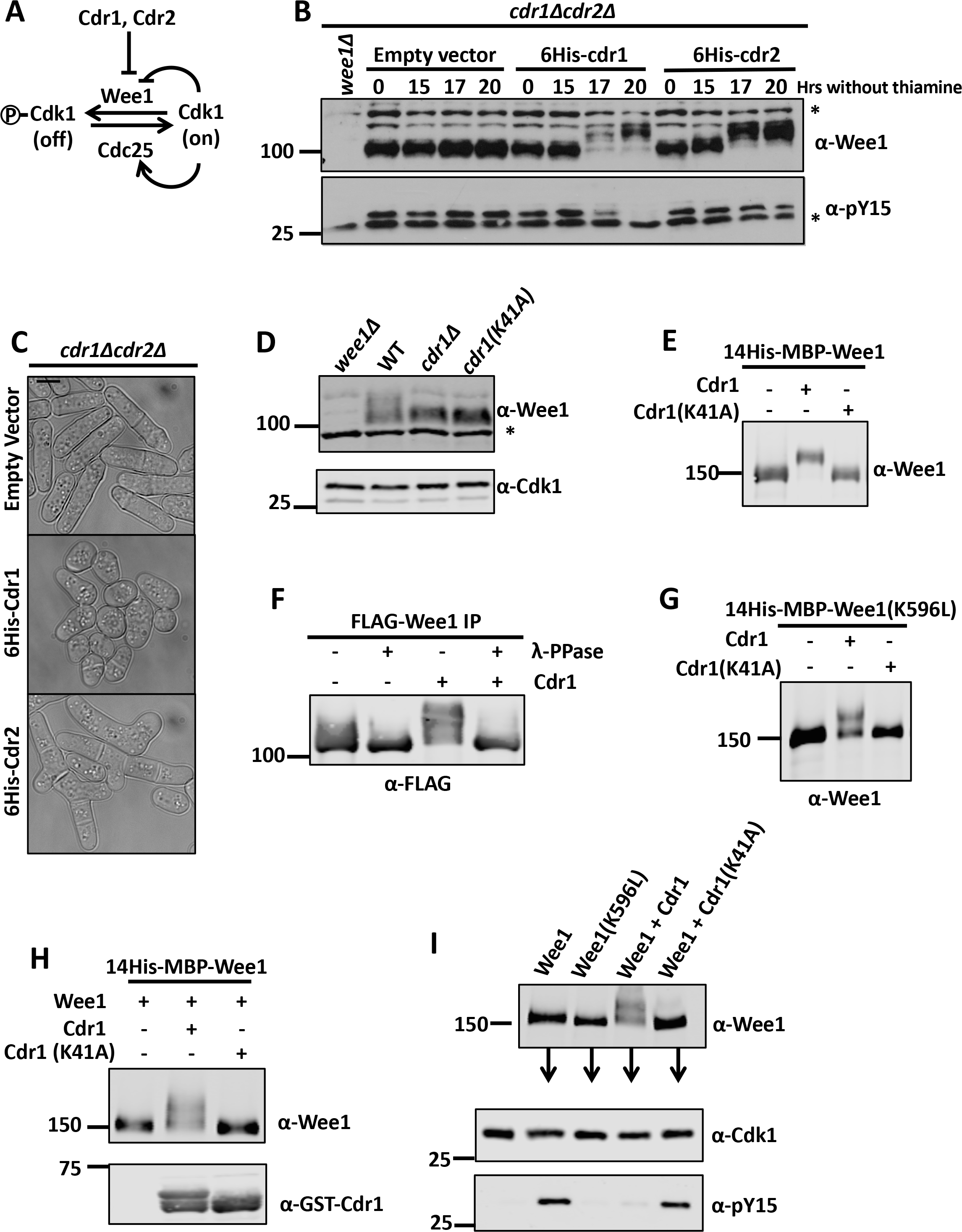
Cdr1 phosphorylates and inhibits Wee1. (A) Schematic of pathway. (B) Whole-cell extracts (WCE) were separated by SDS-PAGE and western blotted against endogenous Wee1. Cdk1-pY15 levels were monitored for Wee1 activity, asterisks mark background bands. (C) DIC images of *cdr1*Δ*cdr2*Δ cells with over-expression plasmids. Scale bar, 5 µm. (D) WCE were separated by SDS-PAGE and blotted against endogenous Wee1. Cdk1 is used as a loading control, asterisk denotes background band. (E) Cdr1 phosphorylates Wee1 in Sf9 cells. Wee1 was co-expressed with Cdr1 or Cdr1(K41A) in Sf9 cells. (F) Cdr1-dependent band shift is due to phosphorylation of Wee1. Wee1 was expressed alone or co-expressed with Cdr1, immunoprecipated, and treated with λ-phosphatase. (G) Co-expression of Wee1(K596L) with Cdr1/Cdr1(K41A) in Sf9 cells. (H) Cdr1 phosphorylates Wee1 directly *in vitro*. GST-Cdr1(1-354) was expressed and purified from bacteria, and mixed with ATP and purified 14His-MBP-Wee1. (I) Cdr1-dependent phosphorylation of Wee1 inhibits Wee1 kinase activity. Wee1 was phosphorylated by Cdr1 as in (H), and then incubated with Cdk1-Cdc13 immunoprecipitated from *S. pombe*. Cdk1-pY15 was used to monitor Wee1 kinase activity.

Previous studies found that Cdr1 can directly phosphorylate and inhibit Wee1 kinase activity *in vitro* (Coleman *et al*., 1993; Parker *et al*., 1993; Wu and Russell, 1993). However, in budding yeast, the Cdr1-like kinase Hsl1 does not phosphorylate or inhibit the Wee1-like kinase Swe1 (Kellogg, 2003; Sakchaisri *et al*., 2004; Keaton and Lew, 2006). These conflicting results indicate that a more detailed analysis of the mechanism for Cdr1-Wee1 regulation is needed. Consistent with previous work (Coleman *et al*., 1993; Parker *et al*., 1993; Wu and Russell, 1993), co-expression of Wee1 and active Cdr1 in Sf9 insect cells caused a phosphorylation-dependent shift in the SDS-PAGE migration of Wee1 (Fig 1E-F). Phosphorylation of Wee1 was not induced by co-expression with catalytically inactive *cdr1(K41A)* (Fig 1E). Further, the shift was not due to auto-phosphorylation because we observed a similar result using the inactive mutant *wee1(K596L)* (Fig 1G). As a more direct test, we performed *in vitro* kinase assays with purified proteins (Fig S1A-E) including the active construct Cdr1(1-354), which was expressed and purified from bacteria. Cdr1 directly phosphorylated Wee1, but Cdr1(K41A) did not (Fig 1H). We performed two-step *in vitro* kinase assays to test the effects of this phosphorylation on Wee1 activity. Wee1 that was phosphorylated by Cdr1 did not phosphorylate its substrate Cdk1-Y15, whereas Wee1 retained activity after incubation with Cdr1(K41A) (Fig 1I). Taken together, our results show that Cdr1 phosphorylates Wee1 in fission yeast cells, insect cells, and *in vitro*. Our findings confirm and extend past work showing that Cdr1 directly phosphorylates Wee1, and this modification inhibits Wee1 kinase activity (Coleman *et al*., 1993; Parker *et al*., 1993; Wu and Russell, 1993).

### Cdr1-dependent phosphorylation sites on Wee1

Next, we sought to identify the sites on Wee1 that are targeted by Cdr1 for inhibitory phosphorylation. We used LC-MS/MS to map Wee1 phosphorylation sites from three independent experiments (Table S2). First, we purified Wee1 from insect cells following its expression either alone or in combination with Cdr1. We identified sites that were phosphorylated specifically upon co-expression with Cdr1. Second, we performed a similar experiment with the inactive mutant Wee1(K596L) to ensure that identified sites were not due to auto-phosphorylation. Third, we performed *in vitro* kinase assays by mixing purified Wee1 or inactive Wee1(K596L) with either active Cdr1 or inactive Cdr1(K41A), and then mapped Wee1 phosphorylation sites specifically induced by active Cdr1. A number of phosphorylation sites were identified in multiple experiments throughout the Wee1 sequence (Fig S2B). We focused primarily on sites in the kinase domain because Cdr1 regulates Wee1 kinase activity and phosphorylates the kinase domain alone (Fig 2A, S2A) (Coleman *et al*., 1993; Parker *et al*., 1993).

**Figure 2:**
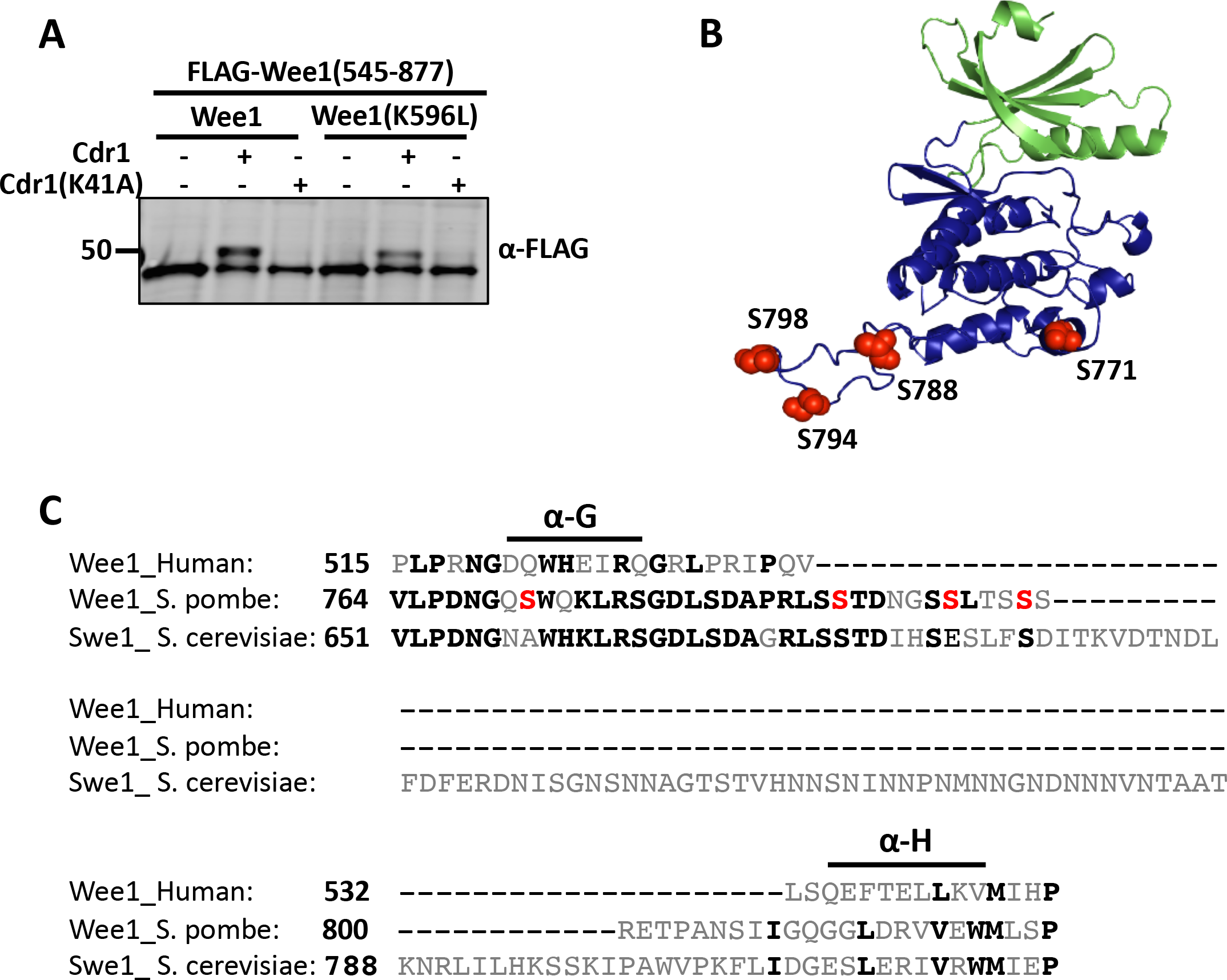
Identification of Cdr1-dependent phosphorylation sites on Wee1. (A) Cdr1 phosphorylates the Wee1 kinase domain. 10His-Cdr1 or 10His-Cdr1(K41A) was co-expressed in Sf9 cells with either FLAG-Wee1(545-877) or FLAG-Wee1(545-877; K596L). (B) Model of *S. pombe* Wee1 kinase domain threaded into human Wee1 from SWISS-MODEL. Green region indicates the N-terminal lobe; blue highlights the C-terminal lobe. Phosphorylated residues in the extended loop are marked in red. (C) Sequence alignment of Wee1 from human, *S. pombe*, and *S. cerevisiae*. Red serines are phosphorylated by Cdr1. Black amino acids are conserved.

To pinpoint which of these phosphorylation sites mediate inhibition of Wee1 by Cdr1 in cells, we generated a panel of mutants in which different phosphorylated residues were changed to alanine, thereby preventing phosphorylation. We reasoned that a non-phosphorylatable Wee1 mutant would be hyperactive, leading to an elongated cell length at division similar to *cdr1*Δ cells. These constructs were integrated into the genome and expressed by the *wee1* promoter as the sole copy in these cells. By analyzing combinations of mutations, we determined that some mutations (*e.g*. S21A and S822A) had no effect on cell size, while others (*e.g*. S781A) caused a loss-of-function *wee* phenotype (Fig S2C and Table S1). Importantly, we generated one mutant that mimicked the *cdr1*Δ phenotype. We named this mutant *wee1(4A)* because it prevents phosphorylation at 4 sites: S771, S788, S794, and S798.

The phosphorylation sites mutated in *wee1(4A)* are clustered within the C-lobe of the kinase domain and have interesting regulatory potential. Using sequence alignments and structural modeling, this cluster falls mostly within a loop that connects α-helices G and H of the Wee1 kinase domain (Fig 2B-C) (Squire *et al*., 2005). This loop is extended in *S. pombe* Wee1 as compared to human Wee1, so the sites are not obviously present in the human polypeptide sequence. Interestingly, this loop is dramatically extended and asparagine-rich in the *S. cerevisiae* Swe1 sequence, and therefore the conserved sites may not be subject to a related regulatory mechanism (Fig 2C). Several eukaryotic protein kinases have extended loops connecting α-G and α-H within the GHI subdomain (Hanks and Hunter, 1995; Scheeff and Bourne, 2005). This subdomain acts as a substrate docking site and connects to the activation segment to regulate catalytic activity (Deminoff *et al*., 2009; Taylor *et al*., 2012). Thus, post-translational modifications in this region have the potential to regulate kinase activity.

### *wee1(4A)* prevents regulation by Cdr1

We performed a series of *in vivo* experiments to test key predictions for the *wee1(4A)* mutant. First, if Cdr1 functions by phosphorylating these residues, then cellular phosphorylation of Wee1 should be reduced in the *wee1(4A)* mutant. Consistent with this model, wee1(4A) phosphorylation was reduced when compared to wild type, and its phosphorylation was not altered by *cdr1*Δ or *cdr2*Δ (Fig 3A). Second, if both *cdr1*Δ and *wee1(4A)* mutants are elongated at division due to the same pathway, then combining these two mutations should not generate additive or synthetic defects. Indeed, *cdr1*Δ *wee1(4A)* cells divided at the same size as *cdr1*Δ cells (Fig 3B and Table S1). Third, *wee1(4A)* mutations should be additive or synthetic with mutations in *cdc25*. Past work has shown that *cdr1* mutations are synthetically lethal with *cdc25* mutant alleles (Young and Fantes, 1987). Both *wee1(4A)* and *cdr1*Δ were synthetically lethal with *cdc25-dD* (Fig 3C). These combined experiments show that *wee1(4A)* recapitulates the *cdr1*Δ phenotype and prevents Wee1 phosphorylation in cells.

**Figure 3:**
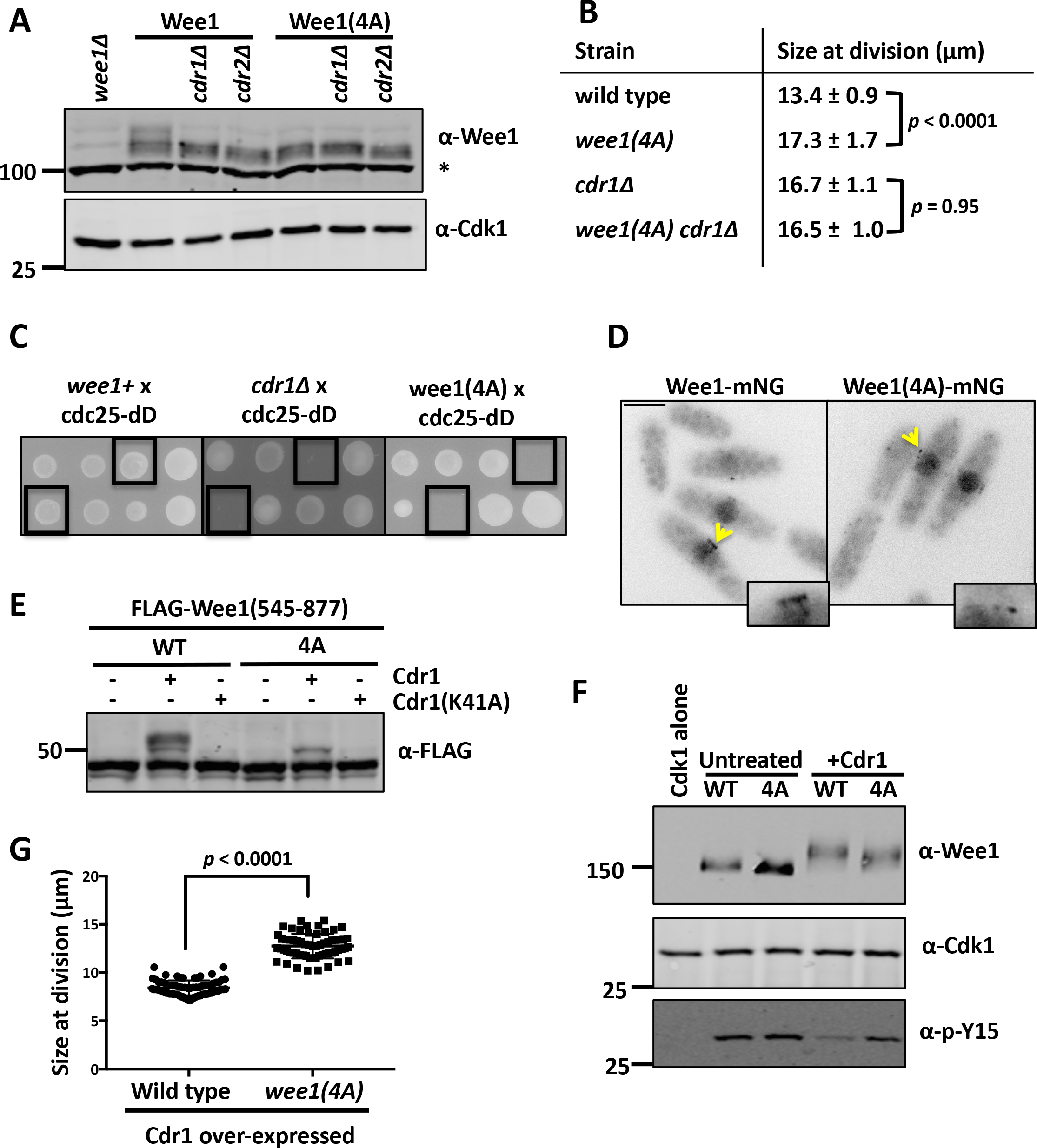
Phenotypes of phosphorylation-resistant *wee1(4A)* mutant. (A) WCE samples were separated by SDS-PAGE, and blotted to detect Wee1. Cdk1 is used as a loading control, asterisk marks background band. (B) Cell size measurements of the indicated strains. Values are mean ± SD for n>50 each strain. *p* value for one-way ANOVA shown for wild type vs *wee1(4A)*, and for *cdr1*Δ vs *wee1(4A) cdr1*Δ. (C) Both *wee1(4A)* and *cdr1*Δ are synthetically lethal with *cdc25-dD*. Black boxes indicate double mutants from tetrad analysis. (D) Localization to nucleus and cortical nodes is unaffected in the *wee1(4A)* mutant. Middle focal plane images. Scale bar, 5 µm. Boxes are enlarged images of nodes denoted by a yellow arrow. (E) *4A* mutation prevents Cdr1-dependent phosphorylation of the Wee1 kinase domain. Co-expression of wild type or Wee1(4A) with Cdr1/Cdr1(K41A). (F) Wee1(4A) has impaired Cdr1 inhibition. Wee1 kinase activity was tested as in Fig 1I. (G) Cell size at division for the indicated strains, showing mean and SD for n>50 cells each. *p* value from unpaired T-test.

Next, we addressed potential mechanisms that could explain the phenotype of *wee1(4A)*. We confirmed that wee1(4A) protein level does not increase and still localizes to cortical nodes (Fig 3A,D). However, Cdr1 was unable to induce hyperphosphorylation of the wee1(4A) kinase domain in insect cells (Fig 3E, S3C). In two-step *in vitro* kinase assays, Cdr1 readily inhibited Wee1 but not wee1(4A) (Fig 3F). Unlike wild type Wee1, the wee1(4A) mutant still phosphorylated Cdk1-pY15 after treatment with Cdr1. These results indicate that the primary defect of the wee1(4A) mutant is loss of inhibition by Cdr1. Accordingly, the size of *wee1(4A)* cells was largely (but not entirely) insensitive to Cdr1 overexpression (Fig 3G). Taken together, our results show that phosphorylation of these four residues by Cdr1 inhibits Wee1 activity. We note that the 4A mutation does not completely abolish Wee1 phosphorylation and regulation by Cdr1 *in vitro*, and Cdr1 phosphorylates additional residues both within and beyond the Wee1 kinase domain. Thus, *wee1(4A*) explains regulation by Cdr1 *in vitro* and *in vivo*, but we do not rule out a role for phosphorylation of additional residues in Cdr1-Wee1 regulation.

Wee1 has also been shown to be phosphorylated and inhibited by Cdk1 in budding yeast, humans, and other systems (Mueller *et al*., 1995; Harvey *et al*., 2005). This feedback sharpens the ultrasensitive nature of Cdk1 activation for mitotic entry (Pomerening *et al*., 2003; Sha *et al*., 2003; Kim and Ferrell, 2007). Whether this feedback occurs in fission yeast and its regulation by Cdr1 has not been tested. We used the analog-sensitive Cdk1-asM17 allele (Aoi *et al*., 2014), which can use bio-orthoganol 6-Bn-ATPγS to thiophosphorylate direct substrates (Allen *et al*., 2007; Hertz *et al*., 2010). We confirmed that *S. pombe* Cdk1-asM17 directly thiophosphorylates Wee1 and Wee1(K596L) (Fig 4A). Thus, Cdk1 phosphorylates Wee1 in fission yeast, similar to other systems. Next, we found that Wee1 phosphorylation by Cdr1 did not impair the level of thiophosphorylation by Cdk1-asM17 (Fig 4B). We conclude that Cdr1 phosphorylation of Wee1 does not block subsequent phosphorylation by Cdk1.

**Figure 4:**
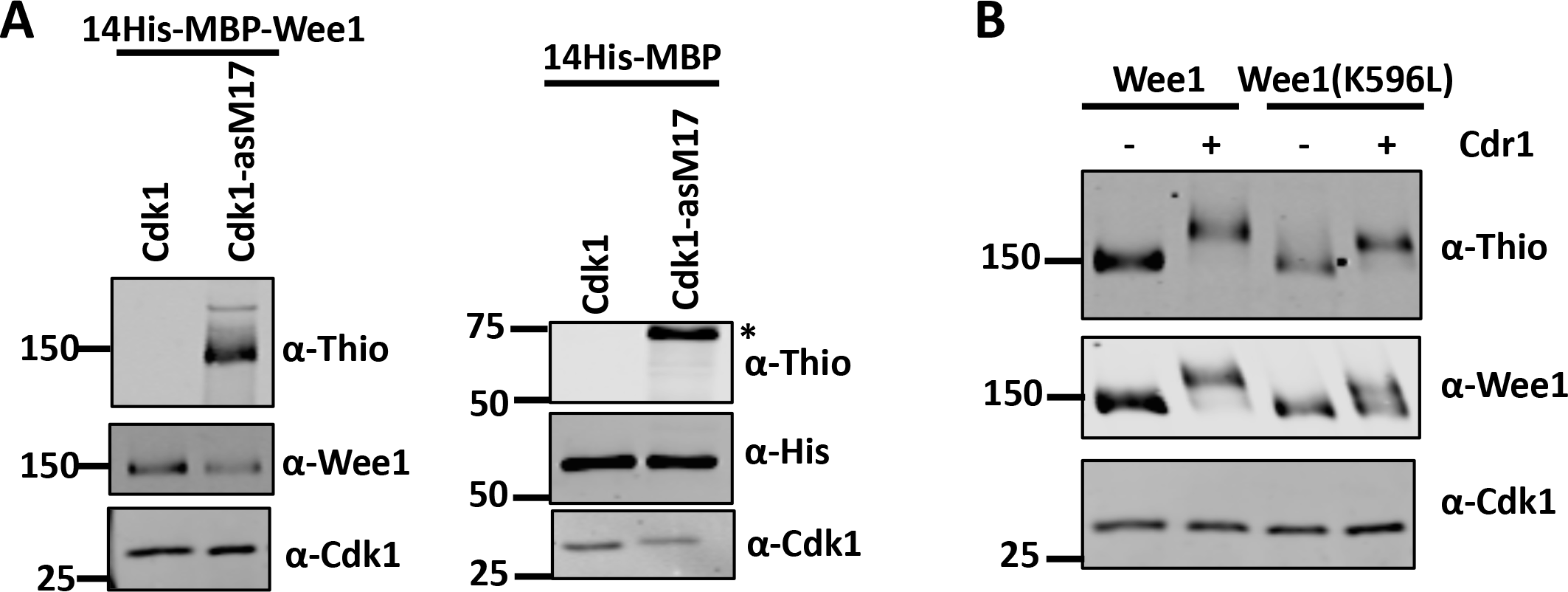
Cdk1 phosphorylation of Wee1 is not blocked by Cdr1. (A) Cdk1 phosphorylates Wee1 *in vitro*. Left: Wild type Cdk1 or Cdk1-asM17 was immunoprecipitated and incubated with purified 14His-MBP-Wee1. These *in vitro* assays used bio-orthoganol 6-Bn-ATPγS, which can only be utilized by Cdk1-asM17 to thiophosphorylate direct substrates. Right: Cdk1-asM17 does not phosphorylate 14His-MBP. (*) denotes background band. (B) Cdk1-asM17 thiophosphorylates both Wee1 and Wee1(K596L). This activity is not blocked by prior phosphorylation of Wee1 constructs by Cdr1.

### Spatial control of Cdr1-Wee1 signaling in cells

Cdr1 and Wee1 both localize to cortical nodes in the cell middle (Martin and Berthelot-Grosjean, 2009; Moseley *et al*., 2009; Allard *et al*., 2018). These nodes are formed by Cdr2 oligomers that are required for Cdr1 and Wee1 recruitment (Martin and Berthelot-Grosjean, 2009; Moseley *et al*., 2009; Allard *et al*., 2018). We previously identified a mutant *cdr1(*Δ*460-482)* that fails to localize to nodes and results in elongated cell size at division like *cdr1*Δ (Fig 5A) (Opalko and Moseley, 2017). We tested the effects of artificially recruiting mEGFP-cdr1(Δ460-482) back to nodes using cdr2-GBP-mCherry, which contains the GFP-binding peptide (GBP). In this system, mEGFP-cdr1(Δ460-482) colocalized with cdr2-GBP-mCherry at nodes (Fig 5B). More importantly, recruitment back to nodes had strong effects on cell size and Wee1 phosphorylation (Fig 5C-D and Table S1). In the *cdr1(*Δ*460-482)* mutant, Wee1 was not hyperphosphorylated and cells divided at an increased size, similar to *cdr1*Δ. However, recruitment of mEGFP-cdr1(Δ460-482) back to nodes by cdr2-GBP-mCherry caused Wee1 to be even more hyperphosphorylated than in wild type cells. A similar effect was seen using full-length mEGFP-cdr1. Along with enhanced Wee1 hyperphosphorylation, these cells divided at a smaller size than wild type cells. These results show that Cdr1 localization to nodes is a limiting factor for phosphorylation of Wee1 and cell size at division. Further, they demonstrate that cdr1(Δ460-482) retains activity but can only phosphorylate Wee1 at nodes, strongly supporting a model where inhibitory phosphorylation of Wee1 occurs at nodes.

**Figure 5:**
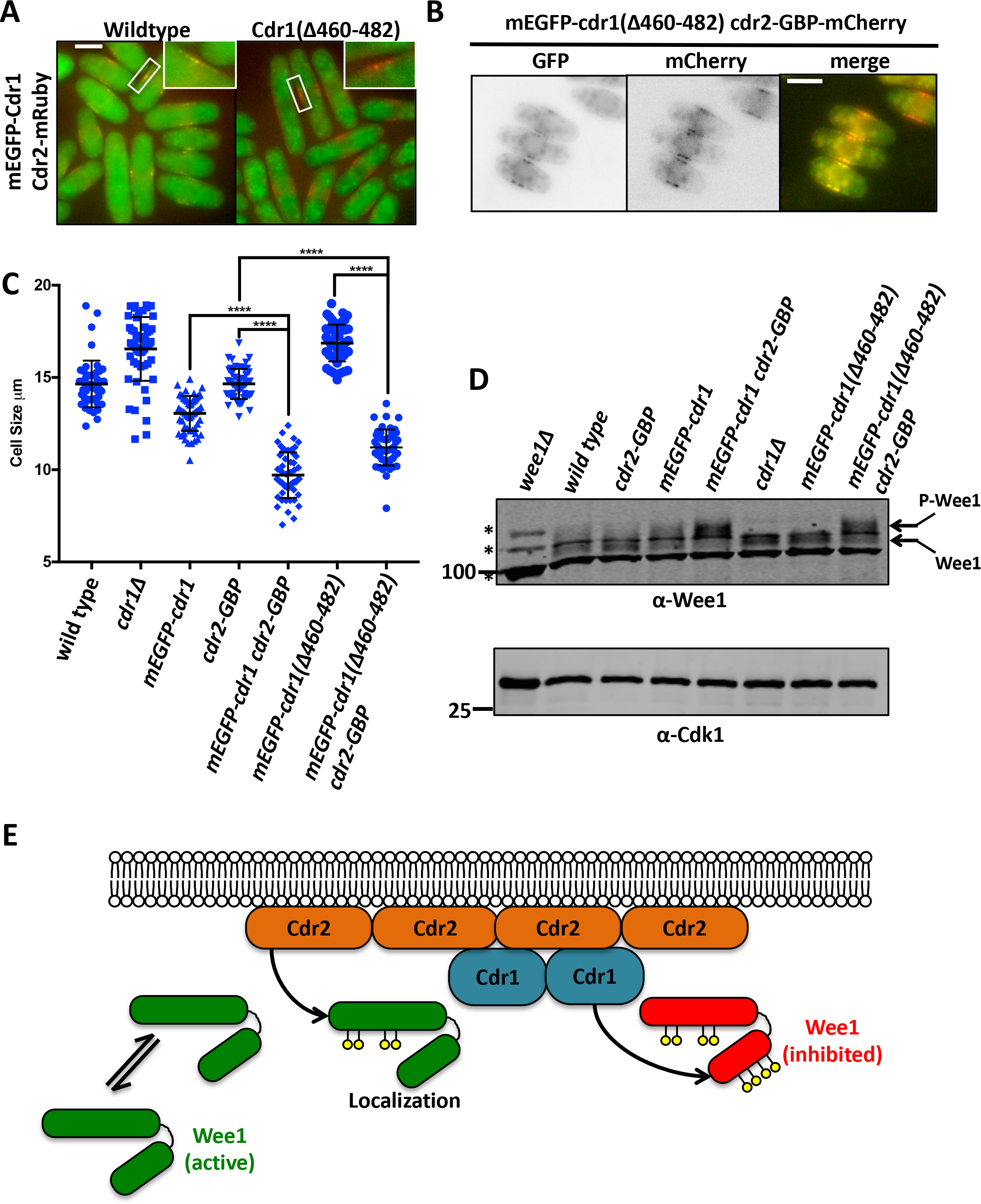
Wee1 inhibition occurs at nodes. (A) mEGFP-tagged Cdr1 or Cdr1(Δ460-482) with Cdr2-mRuby. Scale bar, 5 μm. (B) Colocalization of mEGFP-cdr1(Δ460-482) Cdr2-GBP-mCherry. Scale bar, 5 µm. (C) Tethering Cdr1(Δ460-482) to nodes rescues size defect. Cell size measurements of indicated strains. Line marks mean; error bars mark SD, n>50 for each strain. A one-way ANOVA was used for statistical comparisons. **** indicates *p* < 0.0001. (D) Cdr1 localization dictates Wee1 phosphorylation state. WCEs were separated by SDS-PAGE. Note upper, hyperphosphorylated band for Wee1 upon node targeting of Cdr1. Cdk1 was used as a loading control, asterisk marks background band. (E) Model for two-step inhibition of a Wee1 molecule at a node.

Our results combined with past work suggest a two-step mechanism for regulation of a Wee1 molecule at nodes (Fig 5E). Wee1 localizes to nodes in transient bursts that last between 5-15 seconds (Allard *et al*., 2018). This localization requires Wee1’s non-catalytic N-terminus, which can be phosphorylated by Cdr2 (Kanoh and Russell, 1998; Allard *et al*., 2018). In kinase-dead *cdr2* mutants, Wee1 localizes to nodes in extremely short bursts (< 2 seconds) (Allard *et al*., 2018). As the first step towards inhibition, we propose that Cdr2 phosphorylates the Wee1 N-terminus to slow the off-rate for Wee1 release, thereby “trapping” Wee1 in the node. In the second step, Cdr1 within the node can then phosphorylate the Wee1 kinase domain to inhibit catalytic activity of this molecule through the mechanism described in this study. These two steps are consistent with many past results and show how these two related kinases control distinct aspects of a shared inhibitory mechanism. An open question going forward is why this reaction needs to occur at nodes. It is possible that nodes simply increase the local concentration of Cdr1 and Wee1. Indeed, both Cdr1 and Wee1 are expressed at low levels, which could preclude an efficient interaction in the cytoplasm. This possibility is supported by the finding that Cdr1 overexpression bypasses the need for Cdr2, leading to Wee1 hyperphosphorylation, inhibition of Wee1, and reduced cell size (Fig 1B-C). An alternative possibility is that the structure of a node promotes efficient phosphorylation of Wee1, for example by promoting a more active conformation of Cdr1. These and other possibilities are not mutually exclusive, and additional work on this system may reveal general principles for the emerging theme of signal transduction in cortical clusters.

In conclusion, we have answered long-standing questions regarding how Cdr1/Nim1 inhibits Wee1 in fission yeast cells. Cdr1 inhibits the kinase activity of Wee1 by directly phosphorylating a cluster of residues connecting α-helices G and H. The sequence of this region is evolutionarily divergent, likely explaining why Cdr1-like kinases may not act directly on Wee1 in some other species. However, the functional role of this region within kinase domains suggests that other kinases with insertions at the GHI subdomain may be regulated by mechanisms similar to the one we describe here. Phosphorylation of these residues in Wee1 explains the cell size defects of *cdr1*Δ mutants, and likely represent the functional output of the well-studied Pom1-Cdr2-Cdr1-Wee1 pathway (Martin and Berthelot-Grosjean, 2009; Moseley *et al*., 2009; Pan *et al*., 2014; Allard *et al*., 2018, 2019; Gerganova *et al*., 2019). Efficient signaling in this pathway requires organization within oligomeric clustered structures at the cell cortex. Given the critical role of these structures in relaying cell size information to the core cell cycle machinery, additional insight into the underlying mechanisms will have implications for cell size control. Molecular clusters are a feature of many other signal transduction pathways, so the Cdr2/Cdr1/Wee1 network will also serve as a model for broader mechanistic studies of signal transduction.

## Materials and Methods

### Yeast strains and growth

Standard *S. pombe* media and methods were used (Moreno *et al*., 1991). Strains and plasmids used in this study are listed in supplementary table S1. Sf9 constructs were cloned via restriction digest into Fastbac vector (Thermo Fisher). Mutations in the Wee1 or Cdr1 gene were made by Gibson assembly (QuantaBio), or by site-directed mutagenesis by Quikchange (Stratagene). Wee1 was expressed in a pJK148 plasmid containing wild type Wee1 along with 1000bp each of the Wee1 promoter and terminator. To introduce *wee1* mutations into cells, we deleted one copy of *wee1+* from a wild type diploid strain (JM5334) using PCR-based homologous recombination to generate the heterozygous diploid strain JM5337. Mutant *wee1* alleles in pJK148-based plasmids were then transformed into the *leu1* locus. Diploid cells were then sporulated, and the resulting spores were separated via tetrad dissection. Colonies that grew on leu-plates and were G418 resistant were verified by colony PCR and by western blot analysis.

To test for synthetic lethality with *cdc25* mutants, we used a *cdc25-degron-DAmP::hygro* strain. *wee1*+ (JM5578), *wee1(4A)* (JM5709), and *cdr1*Δ*::kanMX6* (JM483) were crossed to *cdc25-degron DAMP::hygro* (JM5886/JM5887). A full plate of tetrads was analyzed (9-10 tetrads). Crosses with the *wee1(4A)* yielded a synthetic lethality with *cdc25-degron-DAmP(dD)* both with and without a wild type copy of Wee1 at the endogenous locus. When *cdc25-dD* was crossed to a *cdr1*Δ with the *wee1*Δ*::kanMX6 leu1(Pwee1-wee1-Twee1)* background, *cdr1*Δ cells were also synthetically lethal with *cdc25-degron DAMP* both with and without a wild type copy of Wee1 at the endogenous locus. To look specifically at the genetic interaction between Cdr1 and Cdc25 with one copy of wild type Wee1 present, a *cdr1*Δ*::kanMX6* strain in a wild type background was crossed to *cdc25-dD*. For mEGFP-Cdr1(Δ460-482), Cdr1 was mutated in a pJK148 vector containing 970 bp of Cdr1 promoter, mEGFP-Cdr1 and 300 bp of Cdr1 terminator. Each vector was then integrated into the *leu1* locus in *cdr1*Δ*::kanMX6 leu1-32*.

For Cdr1 or Cdr2 overexpression, pREP3x plasmids were transformed into *cdr1::kanMX6 cdr2*Δ*::ura4 ULA-* (JM2070) cells and grown in EMM-leu +thiamine. Cells were then washed vigorously and resuspended in EMM-leu to induce expression of Cdr1 or Cdr2, starting at time point 0. For experiment in Figure 3G, cells were grown at 32°C in EMM4S lacking thiamine for 19 hours before analyzing. GFP-Cdr1 was over-expressed to the same level in both strains as verified by western blot using α-GFP antibodies.

### Sf9 cells Co-expression and Wee1 purification

Sf9 cells were grown in Grace’s media supplemented (Gibco) with 10% heat inactivated FBS, 10µg/mL Gentamicin and 0.25 µg/mL Amphotericin B, at 27°C. Constructs were expressed in Sf9 cells using Bac-to-Bac Baculovirus expression system (Thermo Fisher). Wee1 constructs were co-expressed with Cdr1/Cdr1(K41A) for 3 days. Cells were resuspended in SDS-PAGE sample buffer (65mM Tris pH 6.8, 3% SDS, 10% glycerol, 10% 2-mercaptoethanol, 50mM sodium fluoride, 50mM β–gylcerophosphate, 1mM sodium orthovanadate, complete EDTA-free protease inhibitor tablet (Pierce)), boiled for 5 mins, and the resulting lysates were separated by SDS-PAGE.

For purification of Wee1 from Sf9 cells, 14His-3C-MBP-Wee1 was expressed in Sf9 cells for 3 days. Cells were then harvested and resuspended in lysis buffer (50mM Tris pH 7.5, 150mM NaCl, 75mM sodium fluoride, 75mM β–gylcerophosphate, 1mM PMSF, 10mM imidazole pH 8, complete EDTA-free protease inhibitor tablet (Pierce)). Cells were lysed by French press. Triton X-100 was added to a final concentration of 1%, and glycerol was added to a concentration of 0.5%. Lysates were clarified by centrifuging at 15,000rpm for 15 minutes at 4°C. Lysates were then added to TALON^®^ (TaKaRa) resin and incubated for 1 hour at 4°C. The resin was then washed with wash buffer (50mM Tris pH 7.5, 500mM NaCl, 75mM sodium fluoride, 75mM β–gylcerophosphate, 20mM imidazole pH 8). Wee1 was then eluted using 500mM imidazole pH 8. For purification and mass spec, Wee1 was purified using the MBP tag due to co-expression with 10His-Cdr1. The approach was similar as above, except for the use of an amylose resin (NEB). Wee1 was then resuspended in elution buffer (1% SDS, 15% glycerol, 50mM Tris pH 8.7, 150mM NaCl) and boiled for 5 minutes. The sample was reduced and alkylated prior to separation by SDS-PAGE.

### Lambda Phosphatase

For lambda phosphatase treatment, FLAG-Wee1 was co-expressed with Cdr1-MBP-14His for 3 days. Cells were harvested and lysed in lysis buffer (20mM HEPES pH 7.4, 1mM EDTA, 300mM NaCl, 0.2% Triton X-100, complete EDTA-free protease inhibitor tablet (Pierce), 75mM sodium fluoride, 75mM β–gylcerophosphate, 1mM sodium orthovanadate, 1mM PMSF) by repeated freeze thaw cycles. Lysates were centrifuged at 15,000 rpm at 4°C for 15 minutes. Clarified lysates were incubated with FLAG M2 magnetic beads (Sigma) for 1 hour. Beads were then vigorously washed in lysis buffer lacking phosphatase inhibitors. Samples were split and treated with 800U of lambda phosphatase (NEB) or mock treated at 30°C for 1 hour.

### *In vitro* kinase assay and Western Blots

GST-Cdr1(1-354) and GST-Cdr1(1-354)(K41A) were expressed in BL21 *E. coli* at 16°C. For these assays, Cdr1 was always freshly purified on the same day that it would be used. Cells were lysed in lysis buffer (1xPBS, 300mM NaCl, 1mM DTT, 75mM sodium fluoride, 75mM β– gylcerophosphate, 1mM PMSF) by French press. Triton X-100 was added to a final concentration of 1% and glycerol to a concentration of 0.5%. Lysates were then centrifuged at 15,000 rpm for 20 minutes at 4°C. Clarified lysates were then incubated on gluthathione-agarose (Sigma) for 1 hour at 4°C. Agarose resin was then washed with wash buffer (1xPBS, 500mM NaCl, 1mM DTT, 75mM sodium fluoride, 75mM β–gylcerophosphate) followed by 50mM Tris pH 7.5, 150mM NaCl. GST-Cdr1 was maintained on glutathione-agarose resin for *in vitro* kinase reactions. Briefly, 0.3µg of purified 14His-3C-MBP-Wee1 was incubated with approximately 5-10µg of GST-Cdr1 in kinase buffer (50mM Tris pH 7.5, 10mM MgCl2 1mM DTT, 2mM ATP, 3µM okadaic acid, 20mM glutathione pH 8) shaking, for 1 hour. To test Wee1 ability to phosphorylate Cdk1, 0.1µg Wee1 was incubated with Cdr1 for 10 minutes. Then reactions were spun down and soluble Wee1 was then added to Cdk1-Cdc13 complexes for 15 minutes. Cdk1-Cdc13 was immunoprecipitated from the fission yeast strain, *cdc13-FLAG wee1-50 mik1*Δ*::ura4+* after growth at the non-permissive temperature for 3 hours. Cdc13-FLAG was then immunoprecipitated as described above.

For thiophosphate ester *in vitro* kinase assay, following *in vitro* kinase assay with Cdr1 as described above, ATP-γS (Axxora BLG-B072-05) was added to each reaction at a final concentration of 50 µM. The reaction was allowed to proceed for 15 minutes. The reaction was then quenched with 20mM EDTA and a final concentration of 2.5mM p-nitrobenzyl mesylate (Abcam Biochemicals) was added and incubated for 2 hours at room temperature. Phosphorylation was then probed using thiophosphate ester antibody RabMAb (ab92570) according to manufacturer’s protocol.

For western blots, whole cell extracts of *S. pombe* were made by flash freezing 2 O.D. of cells. Cells were then lysed in 100µl sample buffer (65mM Tris pH 6.8, 3% SDS, 10% glycerol, 10% 2-mercaptoethanol, 50mM sodium fluoride, 50mM β–gylcerophosphate, 1mM sodium orthovanadate, complete EDTA-free protease inhibitor tablet (Pierce)) in a Mini-beadbeater-16 for 2 minutes. Blots were probed with anti-FLAG M2 (Sigma), anti-GST (Covance), anti-Cdr1 (Opalko and Moseley, 2017), anti-Wee1 (Allard *et al*., 2018), anti-cdc2 (Santa Cruz Biotechnologies sc-53217), anti-pY15 (Cell Signaling #9111L), anti-His (Santa Cruz Biotechnologies sc-8036), and anti-thiophosphate ester (Abcam ab92570).

### Mass spectrometry

Gel bands from Sf9 coexpression of Wee1 and Cdr1 and *in vitro* kinase assays were excised and destained overnight at 37°C using 50mM ammonium bicarbonate/50% acetonitrile. The destained gel bands were protease-digested in 50mM ammonium bicarbonate. To maximize sequence coverage, we tested three different proteases digestions: trypsin at 37°C for 16 hours, GluC/LysC at 37°C for 16 hours, or Proteinase K at 37°C for 1 hour. In our initial mass spectrometry experiment, we found that Proteinase K provided the greatest sequence coverage, while trypsin and GluC/LysC were more limited (see Supplemental Table S2, tab 3). For subsequent experiments, we used Proteinase K digestion, and three separate samples were tested in each condition. This approach dramatically increased our sequence coverage of Wee1. Following digestion, peptides were extracted using 5% formic acid/50% acetonitrile and dried.

Peptides were analyzed on a Q-Exactive Plus mass spectrometer (Thermo Fisher Scientific) equipped with an Easy-nLC 1000 (Thermo Fisher Scientific). Raw data were searched using COMET in high-resolution mode (Eng *et al*., 2013) against *S. pombe* sequence database, with appropriate enzyme specificity, and carbamidomethylcysteine as static modification. Oxidized methionine and phosphorylated serine, threonine, and tyrosine were searched as variable modifications. We used a < 1% false discovery rate to filter the resulting peptide spectral matches. Quantification of LC-MS/MS spectra was performed using MassChroQ (Valot *et al*., 2011) with retention time alignment for smart quantification. Peak areas of Wee1 peptides were normalized to total amount of Wee1 in the sample. Probability of phosphorylation site localization was determined by phosphoRS (Taus *et al*., 2011). All Wee1 phosphorylation sites are provided in Supplemental Table S2. For the Wee1(K596L) experiment in Tab 2, we identified 12 sites only in the presence of Cdr1, 10 sites only in the absence of Cdr1, and 6 sites in both conditions. For the wild type Wee1 experiment in Tab 4, we identified 35 sites only in the presence of Cdr1, 12 sites only in the absence of Cdr1, and 18 sites in both conditions. For the *in vitro* kinase assays in Tab 5, we identified 41 sites only in the presence of active Cdr1, 50 sites only in the absence of active Cdr1, and 21 sites in both conditions. Combining all the sites from Tabs 2, 4, and 5, we identified 58 sites only in the presence of active Cdr1, 32 sites only in the absence of active Cdr1, and 47 sites in both conditions.

For phosphorylation sites identified by mass spectrometry, we generated a panel of non-phosphorylatable mutants (Ser to Ala), a subset of which are shown in Figure S2C. We focused primarily on phosphorylated serines in the kinase domain because Cdr1 phosphorylates the Wee1 kinase domain alone, and also preferentially phosphorylates serines (Coleman *et al*., 1993; Parker *et al*., 1993; Wu and Russell, 1993). Our initial mutagenesis targeted single sites, such as Ser771, that were highly phosphorylated in a Cdr1-dependent manner. All mutants were assayed for cell size at division, and most mutants were also tested by western blot for changes in Wee1 phosphorylation and/or protein abundance *in vivo*. We focused on phosphosite mutants that changed cell size at division and Wee1 phospho-state, but did not influence Wee1 protein abundance and were non-additive with *cdr1*Δ. Because *wee1(S771A)* caused a partial increase in cell size but did not abolish Wee1 phosphorylation in cells, we combined this mutant with nearby phosphosites in the predicted loop region, eventually leading to the *wee1(4A)* mutant.

### Microscopy

Microscopy was performed at room temperature with a DeltaVision Imaging System (Applied Precision), equipped with an Olympus IX-71 inverted wide-field microscope, a Photometrics CoolSNAP HQ2 camera, and Insight solid-state illumination unit. All images are single focal plane images with a 5µm scale bar. Blankophor was used to identify septating cells for cell size measurements. ImageJ was used to measure cell length.

### SWISS-MODEL and Sequence Alignment

SWISS-MODEL was used to thread the kinase domain sequence of *S. pombe* Wee1 into the crystal structure of human Wee1 kinase domain (Guex *et al*., 2009). PyMOL Molecular Graphics system, Version 2.0 Schrödinger, LLC was used to visualize the model. For sequence alignments, clustal omega was used to align the Wee1 amino acid sequences from human, and *S. pombe* Wee1 and *S. cerevisiae* Swe1 (Madeira *et al*., 2019).

### Statistics

All graphs and statistical analysis was performed using GraphPad Prism 8.0.2 (GraphPad Software, La Jolla, CA). A one-way ANOVA using Dunnett’s multiple comparison test was used to in figure S2C to compare each mutant strain to the wild type control. A one-way ANOVA with Tukey’s multiple comparison was used to compare more than 2 data sets.

## Acknowledgments

We thank Moseley lab members for comments and feedback; N. Ramirez and A. Grassetti for technical assistance; M. Sato for strains; and the Biomolecular Targeting Core (BioMT) (P20-GM113132) for equipment. This work was supported by grants from the American Cancer Society (RSG-15-140-01) and the NIH/NIGMS (R01GM099774) to J.B.M, and by grants from NIH/NIGMS (R35GM119455, P20GM113132) to A.N.K., and by the NCCC support grant from NIH/NCI (P30CA023108).

## Figure legends

**Supplemental Figure S1:**
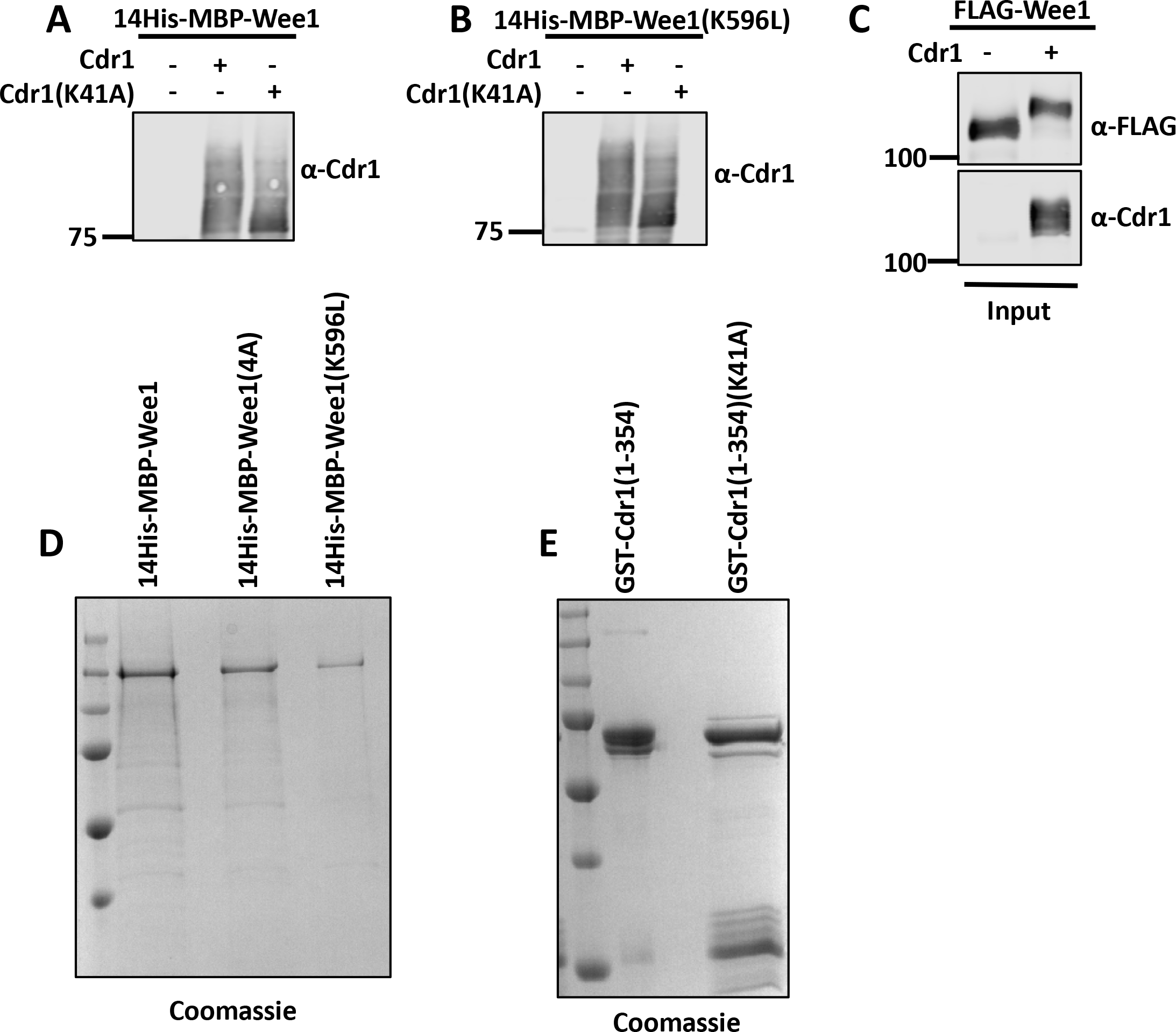
Cdr1 phosphorylates and inhibits Wee1. (A) Western blot for the indicated Cdr1 constructs in Figure 1E. (B) Western blot for the indicated Cdr1 constructs in Figure 1G. (C) Input of Wee1 and Cdr1 from Sf9 cells prior to immunoprecipitation used for Figure 1F. (D) Coomassie-stained SDS-PAGE gel of purified Wee1 constructs used for *in vitro* kinase assays. (E) Coomassie-stained SDS-PAGE gel of purified GST-Cdr1(1-354) and GST-Cdr1(1-354)(K41A) used for *in vitro* kinase assays.

**Supplemental Figure S2:**
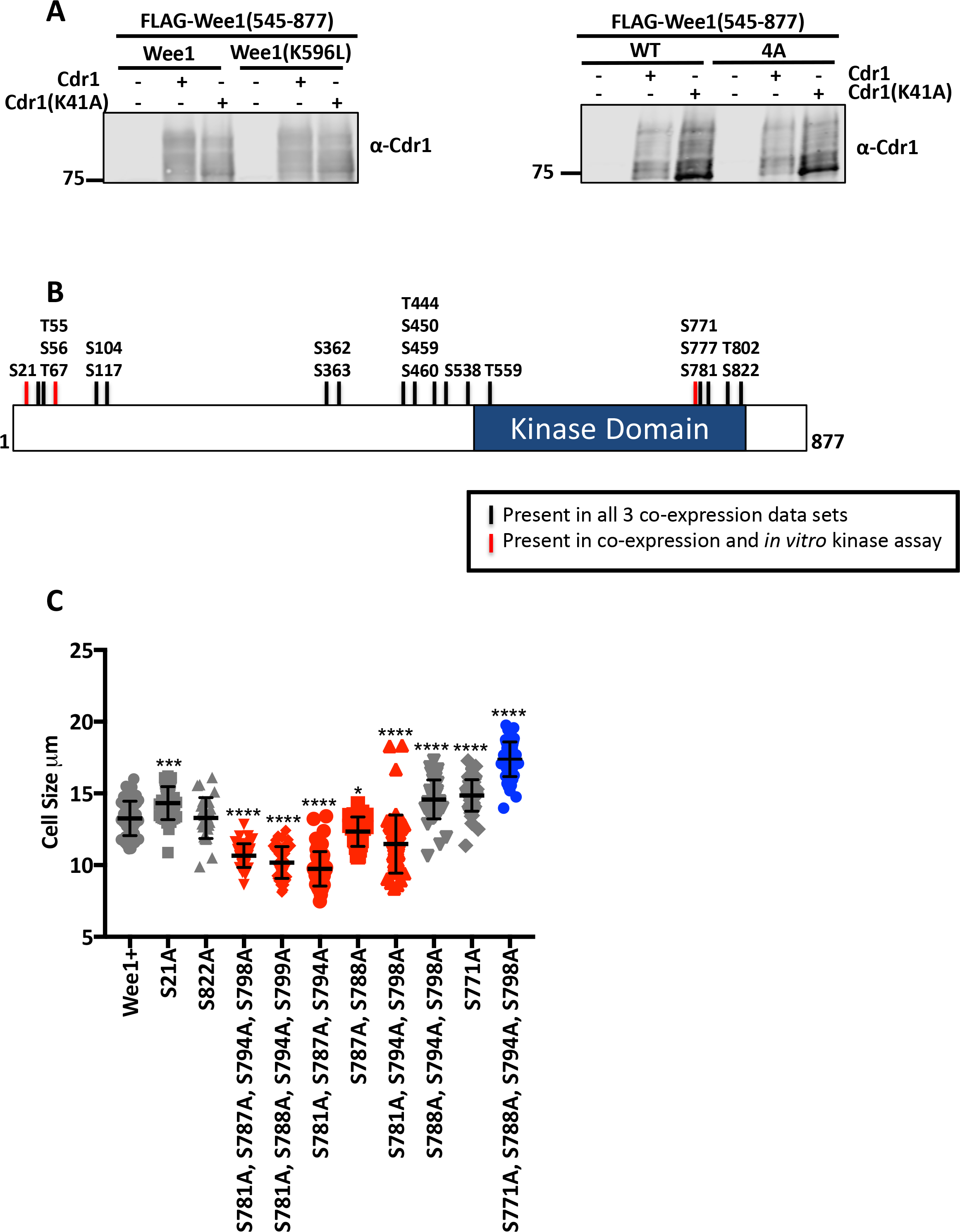
Identification of Cdr1-dependent phosphorylation sites on Wee1. (A) Western blot for Cdr1 expression in the samples analyzed in Figures 2A and 3E. (B) Schematic of Wee1 with phosphorylated residues. Black bars indicate residues that were phosphorylated in all 3 co-expression data sets. Red bars are sites that were present in the co-expression data sets as well as the *in vitro* kinase assay data set. (C) Cell size measurements of serine to alanine mutations. n>25. Grey strains are alleles with either no cell size phenotype or with partial Wee1 gain-of-function phenotype. Red strains indicate loss-of-function of Wee1; *wee1(4A)* is presented in blue. A one-way ANOVA was used to compare each strain to the Wee1+ control. (*) indicates statistical significance. * *p* = 0.0175; *** *p* = 0.0004; **** *p* < 0.0001.

**Supplemental Table S1. Strains, plasmids, and statistics used in this study.**

**Supplemental Table S2. Mass spectrometry data generated in this study.**

