## Supplemental Table S1 for "A mechanism for how Cdr1/Nim1 kinase promotes mitotic entry by inhibiting Wee1"

| Strains | | Source |
|---|---|---|
| JM366 | 972 h- | Lab stock |
| JM483 | nim1Δ::kanMX6 h+ | Lab stock |
| JM777 | wee1Δ::ura4+ ura4-D18 leu1-32 h- | Lab stock |
| JM2070 | nim1Δ::kanMX6 cdr2Δ::ura4+ ULA- | Lab stock |
| JM3474 | cdr1(K41A)h+ | Lab stock |
| JM4957 | wee1-50 mik1Δ::ura4 cdc13-9gly-5 FLAG::kanMX6 leu1-32 ura4-D18 | This study |
| JM5042 | nim1Δ::kanMX6 leu1[Pcdr1-mEGFP-cdr1-Tcdr1] h+ | This study |
| JM5169 | cdr2-GBP-mCherry::hphR h- | This study |
| JM5171 | nim1Δ::kanMX6 leu1[Pcdr1-mEGFP-cdr1(Δ460-482)-Tcdr1] h+ | This study |
| JM5175 | nim1Δ::kanMX6 leu1[Pcdr1-mEGFP-cdr1(Δ460-482)-Tcdr1] cdr2-GBP-mCherry::hphR | This study |
| JM5203 | nim1Δ::kanMX6 leu1[Pcdr1-mEGFP-cdr1-Tcdr1] cdr2-GBP-mCherry::hphR h? | This study |
| JM5256 | nim1Δ::kanMX6 leu1[Pcdr1-mEGFP-cdr1-Tcdr1] cdr2-yomRuby2::hphR | This study |
| JM5257 | nim1Δ::kanMX6 leu1[Pcdr1-mEGFP-cdr1(Δ460-482)-Tcdr1] cdr2-yomRuby2::hphR | This study |
| JM5334 | ade6-M216 ura4-D18 leu1-32/ade6-M210 ura4-D18 leu1-32 h+/h- | This study |
| JM5337 | wee1Δ::kanMX6/wee1+ ade6-M216 ura4-D18 leu1-32/ade6-M210 ura4-D18 leu1-32 h+/h- | This study |
| JM5490 | wee1Δ::kanMX6 leu1[Pwee1-wee1-yomNeonGreen::hphR- Twee1] UA- h+ | This study |
| JM5578 | wee1Δ::kanMX6 leu1[Pwee1-wee1-Twee1] h+ | This study |
| JM5601 | wee1Δ::kanMX6 leu1[Pwee1- wee1(S21A)-Twee1] ura4-D18 A- h? | This study |
| JM5602 | wee1Δ::kanMX6 leu1[Pwee1-wee1(S787A, S788A)-Twee1] ura4-D18 A- h? | This study |
| JM5614 | wee1Δ::kanMX6 leu1[Pwee1- wee1(S781A, S788A, S794A, S799A)-Twee1] ura4-D18 A- h? | This study |
| JM5615 | wee1Δ::kanMX6 leu1[Pwee1-wee1-(S788A, S794A, S798A)-Twee1] ura4-D18 A- h? | This study |
| JM5631 | wee1Δ::kanMX6 leu1[Pwee1- wee1(S771A)-Twee1] ura4-D18 A- h- | This study |
| JM5633 | wee1Δ::kanMX6 leu1[Pwee1- wee1(S822A)-Twee1] ura4-D18 A- h | This study |
| JM5634 | wee1Δ::kanMX6 leu1[Pwee1- wee1(S781A, S787A, S794A)-Twee1] ura4-D18 A- h | This study |
| JM5635 | wee1Δ::kanMX6 leu1[Pwee1- wee1(S781A, S794A, S798A)-Twee1] ura4-D18 A- h | This study |
| JM5658 | cdc2-asM17-bsd leu1-32 ura4-D18 ade6-M210 h90 | MO1893 |
| JM5685 | cdc2-asM17-bsd wee1-50 mik1Δ::ura4 cdc13-9gly-5 FLAG::kanMX6 leu1-32 ura4-D18 | This study |
| JM5686 | wee1Δ::kanMX6 leu1[Pwee1-wee1-Twee1] nim1Δ::natR h? | This study |
| JM5709 | wee1Δ::kanMX6 leu1[Pwee1- wee1(S771A, S788A, S794A, S798A)-Twee1] h- | This study |
| JM5719 | wee1Δ::kanMX6 leu1[Pwee1- wee1(S771A, S788A, S794A, S798A)-Twee1)] nim1Δ::natR h? | This study |
| JM5735 | wee1Δ::kanMX6 leu1[Pwee1- wee1(S771A, S788A, S794A, S798A)-yomNeonGreen::hphR-Twee1] | This study |
| JM5788 | wee1Δ::kanMX6 leu[Pwee1-wee1-Twee1] cdr2Δ::natR h? | This study |
| JM5790 | wee1Δ::kanMX6 leu1[Pwee1- wee1(S771A, S788A, S794A, S798A)-Twee1] cdr2Δ::natR h? | This study |
| JM5867 | cdc25-degron-DAmP::hphR leu1-32 h+ | Lab stock |
| JM6166 | wee1Δ::kanMX6 leu1[Pwee1-wee1(S781A, S787A,S794A, S798A)-Twee1] h? | This study |
| JM6146 | wee1Δ::kanMX6 leu1[Pwee1-wee1-Twee1] KanMx6-P41-nmt-GFP-Cdr1 | This study |
| JM6254 | wee1Δ::kanMX6 leu1[Pwee1- wee1(S771A, S788A, S794A, S798A)-Twee1] KanMx6-P41-nmt-GFP-Cdr1 | This study |

| Plasmids | | Source |
|---|---|---|
| pJM210 | pREP3X | |
| pJM311 | pREP3X-6His-cdr2 | |
| pJM315 | pJK148 | |
| pJM416 | pREP3X-6His-cdr1 | |
| pJM426 | pJK148-Pwee1-wee1+-Twee1 | |
| pJM853 | pJK148-Pcdr1-mEGFP-cdr1-Tcdr1 | |
| pJM885 | pGEX6P1-Cdr1(1-354) | |
| pJM1190 | 14HIS-bd-SUMO-ATG-MBP | pSF1477 Dirk Gorlich |
| pJM1197 | pFastBac-10His-Cdr1 | |

| | |
|---|---|
| pJM1206 | pFastBac-5FLAG-9gly-3C-Wee1 |
| pJM1242 | pFastBac-10His-Cdr1(K41A) |
| pJM1254 | pFastBac-Flag-wee1(545-end) |
| pJM1369 | pFastbac-14his-3c-MBP-Wee1(sf9 codon optimized) |
| pJM1370 | pFastbac-Cdr1-MBP-3c-14his |
| pJM1391 | pFastbac-5Flag-9gly-Wee1(535-end)(K596L) |
| pJM1398 | pJK148-Pcdr1-mEGFP-cdr1(Δ460-482)-Tcdr1 |
| pJM1402 | pFastbac-14his-3C-MBP-Wee1(K596L) sf9 codon optimized |
| pJM1403 | pGEX6P1-Cdr1(1-354)(K41A) |
| pJM1430 | pJK148-Pwee1-wee1(S21A)-Twee1 |
| pJM1432 | pJK148-Pwee1-wee1(S787A, S788A)-Twee1 |
| pJM1435 | pJK148-Pwee1-wee1(S771A)-Twee1 |
| pJM1436 | pJK148-Pwee1-wee1(S781A, S787A,S794A, S798A)-Twee1 |
| pJM1437 | pJK148-Pwee1-wee1(S781A, S788A, S794A, S799A)-Twee1 |
| pJM1438 | pJK148-Pwee1-wee1(S788A, S794A, S798A)-Twee1 |
| pJM1440 | pJK148-Pwee1-wee1(S822A)-Twee1 |
| pJM1441 | pJK148-Pwee1-wee1(S781A, S794A, S798A)-Twee1 |
| pJM1442 | pJK148-Pwee1-wee1(S781A, S787A, S794A)-Twee1 |
| pJM1445 | pJK148-Pwee1-wee1(S771A, S788A, S794A, S798A)-Twee1 |
| pJM1462 | pfastbac-5FLAG-9gly-Wee1(S771A, S788A, S794A, S798A)(545-end) |
| pJM1466 | pfastbac-14HIS-3c-MBP-Wee1(S771A, S788A, S794A, S798A) |

**Stats for Fig 3B**

| **Tukey's multiple comparisons test** | **Significant?** | **Summary** | **Adjusted P Value** |
|---|---|---|---|
| WT vs. *cdr1Δ* | Yes | **** | <0.0001 |
| WT vs. *wee1(4A)* | Yes | **** | <0.0001 |
| WT vs. *wee1(4A)cdr1*Δ | Yes | **** | <0.0001 |
| *cdr1Δ* vs. *wee1(4A)* | Yes | * | 0.0494 |
| *cdr1Δ* vs. *wee1(4A)cdr1*Δ | No | ns | 0.9539 |
| *wee1(4A)* vs. *wee1(4A )cdr1*Δ | Yes | * | 0.0106 |

**Stats for Fig 5C**

| **Tukey's multiple comparisons test** | **Significant?** | **Summary** | **Adjusted P Value** |
|---|---|---|---|
| Wt vs. *cdr1*Δ | Yes | **** | <0.0001 |
| Wt vs. mEGFP-Cdr1 | Yes | **** | <0.0001 |
| Wt vs. Cdr2-GBP-mCherry | No | ns | >0.9999 |
| Wt vs. mEGFP-Cdr1 Cdr2-GBP-mCherry | Yes | **** | <0.0001 |
| Wt vs. mEGFP-Cdr1(Δ460-482) | Yes | **** | <0.0001 |
| Wt vs. mEGFP-Cdr1(Δ460-482) Cdr2-GBP-mCherry | Yes | **** | <0.0001 |
| *cdr1*Δ vs. mEGFP-Cdr1 | Yes | **** | <0.0001 |
| *cdr1*Δ vs. Cdr2-GBP-mCherry | Yes | **** | <0.0001 |
| *cdr1*Δ vs. mEGFP-Cdr1 Cdr2-GBP-mCherry | Yes | **** | <0.0001 |
| *cdr1*Δ vs. mEGFP-Cdr1(Δ460-482) | No | ns | 0.7844 |
| *cdr1*Δ vs. mEGFP-Cdr1(Δ460-482) Cdr2-GBP-mCherry | Yes | **** | <0.0001 |
| mEGFP-Cdr1 vs. Cdr2-GBP-mCherry | Yes | **** | <0.0001 |
| mEGFP-Cdr1 vs. mEGFP-Cdr1 Cdr2-GBP-mCherry | Yes | **** | <0.0001 |
| mEGFP-Cdr1 vs. mEGFP-Cdr1(Δ460-482) | Yes | **** | <0.0001 |
| mEGFP-Cdr1 vs. mEGFP-Cdr1(Δ460-482) Cdr2-GBP-mCherry | Yes | **** | <0.0001 |
| Cdr2-GBP-mCherry vs. mEGFP-Cdr1 Cdr2-GBP-mCherry | Yes | **** | <0.0001 |
| Cdr2-GBP-mCherry vs. mEGFP-Cdr1(Δ460-482) | Yes | **** | <0.0001 |
| Cdr2-GBP-mCherry vs. mEGFP-Cdr1(Δ460-482) Cdr2-GBP-mCherry | Yes | **** | <0.0001 |
| mEGFP-Cdr1 Cdr2-GBP-mCherry vs. mEGFP-Cdr1(Δ460-482) | Yes | **** | <0.0001 |
| mEGFP-Cdr1 Cdr2-GBP-mCherry vs. mEGFP-Cdr1(Δ460-482) Cdr2-GBP-mCherry | Yes | **** | <0.0001 |
| mEGFP-Cdr1(Δ460-482) vs. mEGFP-Cdr1(Δ460-482) Cdr2-GBP-mCherry | Yes | **** | <0.0001 |

**Table for Fig S2C**

| Dunnett's multiple comparisons test | Significant? | Summary | Adjusted P Value |
|---|---|---|---|
| Wee1+ vs. S21A | Yes | *** | 0.0004 |
| Wee1+ vs. S822A | No | ns | 0.9999 |
| Wee1+ vs. S781A, S787A, S794A, S798A | Yes | **** | <0.0001 |
| Wee1+ vs. S781A, S788A, S794A, S799A | Yes | **** | <0.0001 |
| Wee1+ vs. S781A, S787A, S794A | Yes | **** | <0.0001 |
| Wee1+ vs. S787A, S788A | Yes | * | 0.0175 |
| Wee1+ vs. S781A, S794A, S798A | Yes | **** | <0.0001 |
| Wee1+ vs. S788A, S794A, S798A | Yes | **** | <0.0001 |
| Wee1+ vs. S771A | Yes | **** | <0.0001 |
| Wee1+ vs. S771A, S788A, S794A, S798A | Yes | **** | <0.0001 |
